# SafeMut: UMI-aware variant simulator incorporating allele-fraction overdispersion in read editing

**DOI:** 10.1101/2023.03.14.532524

**Authors:** Xiaofei Zhao, Jingyu Guo, Sizhen Wang

**Affiliations:** Genetron Health (Beijing) Co. Ltd, Beijing 102208, China

## Abstract

Next-generation sequencing (NGS) has been widely used for calling biological variants. The gold-standard methodology for accessing the ability of a computational method to call a specific variant is to perform NGS wet-lab experiments on samples known to harbor this variant. Nevertheless, wet-lab experiments are both labor-intensive and time-consuming, and rare variants may not be present in a sample of population. Moreover, these two issues are exacerbated in SafeSeqS which enabled liquid biopsy and minimum-residual disease (MRD) detection with cell-free DNA by using unique molecular identifier (UMI) to detect and/or correct NGS error. Hence, we developed the first UMI-aware NGS small-variant simulator named SafeMut which also considered the overdispersion of allele fraction. We used the tumor-normal paired sequencing runs from the SEQC2 somatic reference sets and cell-free DNA data sets to assess the performance of BamSurgeon, VarBen, and SafeMut. We observed that, unlike BamSurgeon and VarBen, the allele-fraction distribution of the variants simulated by SafeMut closely resembles such distribution generated by technical replicates of wet-lab experiments. SafeMut is able to provide accurate simulation of small variants in NGS data, thereby helping with the assessment of the ability to call these variants in a bioinformatics pipeline.

## Introduction

Nowadays, next-generation sequencing (NGS) is widely used. NGS is especially used in clinical settings to detect cancer mutations. To ensure that we can reliably detect a given mutation, the gold-standard method is to perform multiple wet-lab NGS experiments. However, such method is costly, labor-intensive and time-consuming. Thus, computational methods have been developed to simulate variants in NGS data. The first class of methods, such as ART [1], simulates NGS reads from only the human reference genome without using any pre-existing NGS data. Such methods require neither biological samples nor wet-lab experiments but neither accurately model biological variation nor consider experiment-specific error profiles [2, 3]. Systematic reviews of the first class of methods have been provided [4, 5]. The second class of methods, such as the one used to evaluate the somatic variant caller Mutect [6], mixes *in silico* part of tumor with part of its corresponding paired normal to model tumor with lower variant allele fraction (VAF). Such methods can model experiment-specific error profiles and are trivial to implement, but the repertoire of spiked-in mutations is limited to examples detected previously, and thus already known to be discoverable [2, 3]. Additionally, biological variations, such as copy-number variation [7] and variation in the quantity of originally sampled DNA molecules [8], intrinsically exist in tumors. These variations unfortunately hinder the determination of the expected ground-truth allele fractions to simulate. The third class of methods, such as BamSurgeon and VarBen [2, 3], spikes-in any user-defined mutations to pre-existing sequencing data. Such methods can model both biological variation and experiment-specific error profiles. However, biological variation is also affected by some experimental errors which are not modeled.

Moreover, some new wet-lab methods have been developed to reduce NGS errors. In the Safe Sequencing System (SafeSeqS), molecular barcodes, also known as UMIs (unique molecular identifiers), are attached to the original cell-free DNA (cfDNA) molecules in the preparation of libraries, and the libraries are then sequenced at high depth [9]. Thus, we can observe a mutation in cancer cells as a variant supported by UMI families, where each UMI-family is almost entirely made of fragments supporting only this variant [9]. A low quantity of the DNA released by cancer cells is present in cfDNA in the form of circulating tumor DNA (ctDNA), so SafeSeqS has been widely used in liquid biopsies based on ctDNA assays [10]. Additionally, SafeSeqS has been used in the detection of minimum residual disease (MRD), which is essential for providing adequate treatment to cancer patients [11]. Nowadays, UMI-Gen, which belongs to the first class of methods, is the only UMI-aware NGS-variant simulator [12], so the second and third classes of UMI-aware NGS variant simulators are still lacking. Additionally, duplex sequencing, which attaches two UMIs to each original DNA molecule, can suppress NGS errors even further [13]. Hence, duplex sequencing has numerous applications [14–16]. However, there is no duplex-aware NGS variant simulator yet.

Furthermore, the methodology for evaluating NGS variant simulators has not been well established. Methods belonging to the first class directly simulate NGS data. Thus, researchers compared real NGS data with simulated NGS data in terms of quality-control (QC) metrics such as genome coverage and GC bias [17]. This QC-metrics comparison measures the resemblance of simulated data to real data but does not measure the resemblance of simulated variants to real variants observed in the data. Some researchers compared simulated data and real data in terms of correlation with gold-standard VAFs [3]. Nonetheless, VAFs from simulated data may be better correlated with gold-standard VAFs because simulation may introduce less noise than real NGS experiments, and the correlation between observed and expected values does not measure the similarity between the observed and expected distributions generating these values.

Thus, we proposed the application of a hash-based pseudorandom number generator (PRNG) on UMIs to enable UMI-aware simulation of variants and the use of a log-normal distribution to model the overdispersion observed in the fluctuation of VAFs. We also proposed the methodology of using scatter plots of the mean and variance and of using Q-Q plots of the z-scores to measure the similarity, observed from sequencing data and in terms of VAF, between variants derived from *in silico* simulation and variants derived from wet-lab experiments.

Importantly, we developed an NGS variant simulator named SafeMut. In short, SafeMut spikes-in any user-specified small variant (i.e., SNV and Indel) from a given VCF file into a BAM file which is not supposed to contain this variant. Thus, according to the before-mentioned classification, SafeMut belongs to the third class of methods. Remarkably, the distribution of VAFs simulated by SafeMut is approximately identical to the distribution of VAFs observed from real sequencing data. And SafeMut is aware of the UMIs, including duplex UMIs, in its input BAM file so that each simulated variant is spiked-in to UMI families instead of NGS fragments. Also, the running time and random-access memory usages of SafeMut are similar to that of the fast and memory-efficient command “`samtools view`” which is ubiquitously used in NGS to process SAM/BAM files [18], so SafeMut runs fast and is efficient in terms of its use of computational resources.

## Approach

In brief, we discovered two key insights that can enable the editing of aligned reads, with UMI-awareness and allele-fraction overdispersion, in only one pass over an input BAM file. Our key insights are depicted in (Fig. 1).

**Figure 1:**
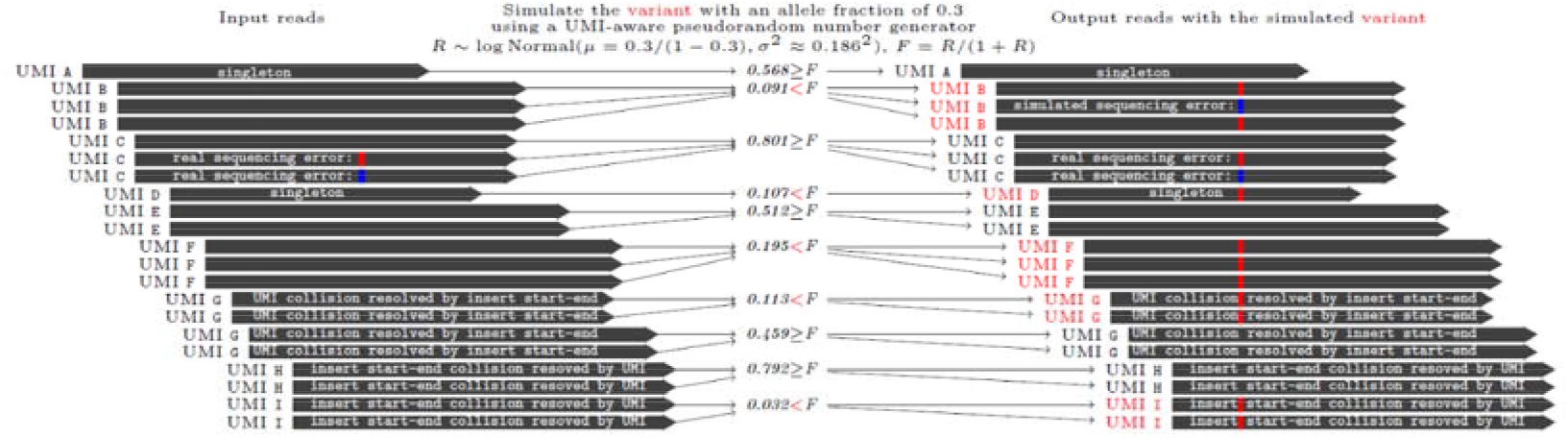
Pseudo-random generation of mutations by SafeMut using unique molecular identifier (UMI)-aware *in silico* modification of reads and incorporating the overdispersion of allele fractions.

First, if a pseudorandom number generator (PRNG) uses the same seed to generate multiple series of pseudorandom numbers, then these series are identical to each other. In a typical NGS dataset, reads derived from the same original DNA molecule have the same start position, end position and UMI tag. Hence, if we derive randomness by using a combination of start position, end position and UMI tag of reads, then reads derived from the same original DNA molecule should be associated with the same random number that can be normalized between zero and one. In this way, we can spike-in a mutation to all these reads grouped together so that the variant occurs at the level of the original DNA molecules. Thereby, PRNG using UMI as its seed can enable UMI-aware simulation of variant.

Second, allele ratio (the number of fragments A supporting the ALT mutant variant allele, divided by the number of fragments B supporting all non-ALT alleles) is known to follow a log-normal distribution [19]. Additionally, the event A/B = 1 is known to be 10^(15/10)^ times more likely than the event A/B = 2 [19]. Hence, we used such lognormal distribution to simulate the overdispersion in the fluctuation of variant allele fractions (VAFs).

The two above-mentioned techniques both require only one pass over the reads found in the input BAM file to simulate all given variants. Hence, our algorithm is supposed to be fast and memory efficient.

## Algorithm

Our algorithm consists of the following main steps.

1. For each read in the sorted input BAM file:

a. Iterate through the input VCF file to discard variants before the start position of this read.
b. Select all remaining variants that overlap with this read.
c. **Compute the hash value H made of the UMI sequence, insert start position and insert end position.** Compute H modulo 2^24^, then divide by 2^24^ to get a pseudorandom real number U between zero and one. If this read is not associated with any UMI, then use the read query name as the UMI sequence. The hash value H is subsequently used as the seed for generating random variable.
d. For each variant that overlaps with this read:

i. **Multiply the allele ratio R by a lognormal random variable** V~logN(μ,σ^2^) generated by using H as the seed to obtain R’, where R = F/(l – F), F is the theoretical variant allele fraction (VAF) of this simulated variant specified in the input VCF file, μ = 0, and where *σ* is set such that the likelihood that V = 2 × exp (μ), divided by the likelihood that V = exp(μ), is 11^-15/10^. Then, let the new VAF F’ be R’/(l+R’).
ii. **If U** < *F′*, **then spike-in the variant into the read.** Otherwise, do not do anything.
e. Output the read with and/or without the spike-in variant(s) into FASTQ files(s).

Allele ratio is defined as the number of fragments supporting the ALT allele divided by the number of fragments supporting all non-ALT (including REF) alleles. Each above step emphasized with bold font is described in more detail as follows.

### Compute the hash value H made of the UMI sequence, insert start position and insert end position

The hash value H of the information that uniquely identifies the original molecule is computed as follows.

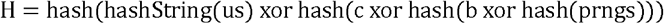

where us is the UMI sequence in string format, b is the insert begin position, c is the insert end position, prngs is a pseudorandom number generator (PRNG) seed, and the operation xor is exclusive or.

Different reads with the same UMI, insert start position and insert end position all have the same hash value H and thus the same pseudorandom number generated for them. Hence, different reads derived from the same original DNA molecule should be either all mutated to the same allele or all not mutated at all if such molecule is observed in sequencing data. Therefore, each simulated variant adds mutation to the original molecules inferred from the sequenced reads, which ensures that reads derived from the same original molecule inferred from sequencing data all represent the same allele at each locus.

### Multiply the allele ratio R by a lognormal random variable

The input VCF file specifies the VAF F to simulate for each variant. In sum, our algorithm introduces some multiplicative stochastic error into F to obtain F’ such that the allele ratio follows a log-normal distribution, where the event that *R*_1_/*R*_2_ = 1 is 10(^15/10)^ times as likely as the event that *R*_1_/*R*_2_ = 2 given two allele ratios *R*_1_ and *R*_2_. This log-normal distribution has been discovered and used in the literature [19, 22].

More specifically, the VAF F is used to calculate allele ratio R as follows: R = F/(l – F). Then, the allele ratio R is multiplied by a random variable V which is distributed as logN(μ = 0, *σ*^2^). The parameter σ is chosen so that if *V*_1_ and *V*_2_ are distributed as logN(μ = 0, *σ*^2^), then the likelihood that *V*_1_/*V*_2_ = 2, divided by the likelihood that *V*_1_/*V*_2_ = 1, is 10^(15/10)^ which is the default value of the -q command-line parameter. If the parameter -q is set to be smaller than zero, then the random variable V always generates one, so the log-normal transformation is disabled in this case.

Next, we generate two standard normal distributions *U*_1_, *U*_2_ ~ N(0,1). Let X be defined as follows.

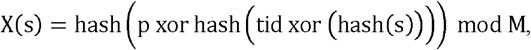

where X is the random variable of interest, p is the genomic position of the variant, tid is the ID of the reference sequence defined in the VCF file format specification [23], s is the seed used to generate X, and M is the maximum possible hash value so that X is normalized to be a real number between zero and one after modulo by M. The function hash has been previously defined. Then, we generate uniformly at random two integers: we let *U*_1_ = *X*(*s*_1_) and *U*_2_ = *X*(*s*_2_), where *s*_1_ and *s*_2_ are by default the sum of the first 50 and 500 hash values of the read names in the input BAM file, respectively. The values 50 and 500 have been arbitrarily selected to introduce randomness into *X*. Then, we use Box-Muller transform to generate a standard normal random variable *Z* from *U*_1_ and *U*_2_ [24].

Next, the lognormal random variable L~exp(σ × Z) is used as the multiplier for the allele ratio R so that R’=L*R is the new allele odds ratio, where σ×Z is equivalent to N(O, σ^2^). Next, we compute the new VAF F’ as follows: F’ = R’/(l + R’)·

### If U < *F*′, then spike-in the variant into the read

If the variant is a single-nucleotide variant (SNV), then we transform the SNV base-call quality (BQ) into its corresponding sequencing-error probability. Next, we generate a pseudorandom number BQN between zero and one as follows.

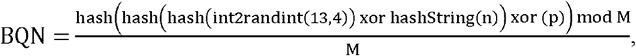

where n is the read name, p is the read start position, and M is the maximum hash value to ensure that BQN is between zero and one. If BQN is less than the sequencing-error probability determined from its corresponding BQ, then we change the original base supporting the SNV into a random base among A, C, G and T. Otherwise, we keep the original base.

If the variant is an insertion/deletion and U < *F*′, then the corresponding bases are always inserted/deleted because insertion/deletion does not have any BQ so that the variant is always spiked-in. The default BQ of each inserted base is 30.

## Implementation

The full implementation of SafeMut is at https://github.com/genetronhealth/safesim/. SafeMut is implemented with C++ using htslib version 1.11. SafeMut accepts one BAM file and one VCF file as input and generates two sets of FASTQ files as output: tumor and normal FASTQ sets containing tumor and normal reads, respectively, to enable the user to identify the tumor/normal origin of each read. The fastq-sort utility from fastq-tools (https://github.com/dcjones/fastq-tools) is then used to sort each output FASTQ file in the two FASTQ sets. Then, the tumor and normal FASTQ sets are merged into one set of FASTQ files. Here, tumor/normal denote any biomaterials that are/aren’t characterized by mutations of interest, respectively. For example, tumor/normal can also denote samples with/without some *de novo* germline mutations, respectively. If the -q command-line parameter is set to a negative number, then the aforementioned log-normal transformation of allele odds ratio is disabled.

## Results

### Overall evaluation procedure

We used two datasets published by the Food-and-Drug Administration (FDA)-led Sequencing Quality Control phase 2 (SEQC2) consortium. These two datasets were produced by the whole-exome sequencing (WES) of the HCC1395 and HCC1395BL cell lines [25] and by the cell-free DNA (cfDNA) sequencing of a well characterized sample [26], respectively (Table S1). These two datasets are both rigorously produced using detailed protocols and open to the general public allowing for permissive usage. More importantly, each of these two datasets is generated by multiple tumor-normal paired sequencing runs and both tumor and normal samples are sequenced at approximately the same depth. Thus, we let each simulator spike-in mutations found in the tumor sample, as specified in the gold-standard tumor variant VCF file, into its matched normal BAM file, to simulate the corresponding tumor BAM. Then, we compared the distributions of variant allele fractions (VAFs) in the tumor BAM files with the VAFs in the simulated BAM files. More specifically, for each variant locus, we first compared the mean and variance of the tumor VAFs with the simulated VAFs. Then, we compared the z-scores, which were normalized by mean and variance, of the tumor VAFs and simulated VAFs. More specifically, we ran the “`bcftools mpileup`” command and then used the FORMAT/AD tag to extract allele depths to compute VAFs. The “`bcftools mpileup`” command already filtered variant read support by mapping quality (MQ), base quality (BQ) and base alignment quality (BAQ) in its computation of allele depths, so our evaluation also indirectly compared simulated variants with real variants in terms of MQ, BQ and BAQ.

We used samtools version 1.11 [27], bcftools version 1.11 [18], BWA MEM version 0.7.17 [28], Picard version 2.26.10 (http://broadinstitute.github.io/picard/), Velvet version 1.2.10 [29], Exonerate version 2.4.0 [30], Fgbio version 1.5.0 (https://github.com/fulcrum-genomics/fgbio), BamSurgeon version 1.3 [2], VarBen commit 0f66e35 [3], and SafeMut version 0.1.5.0b4d0f8 in our evaluation.

We ran all software packages with their default parameters except that we initialized the seeds of the pseudo-random number generators (PRNGs) to zero for BamSurgeon and VarBen. By default, BamSurgeon and VarBen seed their pseudo-random number generators (PRNGs) with the current system time, which leads to the loss of exact reproducibility of variant-simulation results. Hence, we let BamSurgeon and VarBen seed their PRNGs with zero to ensure exact reproducibility without compromising our evaluation.

### Dataset-specific evaluation procedure

For the WES dataset, we used BWA MEM to align all reads to the GRCh38 human reference genome because variant coordinates in the gold-standard VCF are based on GRCh38 [25]. Then, we ran BamSurgeon, VarBen, and SafeMut to spike-in mutations into the normal BAM files which have approximately the same depth as the corresponding tumor BAM files, generating simulated tumor-like BAM files. Next, we compared the VAF distribution of the real tumor BAM files with respect to the VAF distribution of the simulated tumor-like BAM files. A few variants have exactly zero reads supporting the variant allele. Thus, every variant having zero read support for the variant allele in any sample is discarded in our evaluation to prevent the issue of division by zero.

For the cfDNA dataset, we only used the data submitted by Illumina (ILM) and IDT, so we did not use the data submitted by ROC, BRP and TFS because ROC only provided reads processed by UMI-consensus, BRP did not provide UMI information, and TFS data generated runtime errors for both BamSurgeon and VarBen. For the cfDNA dataset, we used BWA MEM to align all reads to the HS37D5 human reference genome because variant coordinates in the gold-standard VCF are based on HG19/HS37D5 [31]. Then, we assumed that all simulators are UMI-aware and thus used the simulation-then-consensus evaluation strategy. In this strategy, we first ran BamSurgeon, VarBen and SafeMut to spike-in mutations into the normal BAM files which have approximately the same depth as the tumor BAM files, generating simulated tumor-like BAM files. Then, we used Fgbio to collapse the sequenced reads into duplex consensus sequences (DCSs) representing the double-stranded original DNA molecules. Next, we compared the DCS VAF distribution of the real tumor BAM files with respect to the DCS VAF distribution of the simulated tumor-like BAM files.

The aforementioned simulation-then-consensus strategy provides comprehensive evaluation for UMI-aware variant simulators. However, BamSurgeon and VarBen are not UMI-aware, so we also used the consensus-then-simulation strategy as a work-around to let UMI-unaware simulators incorporate the effect of the labeling of reads by UMIs in their simulations. In the consensus-then-simulation strategy, we first used Fgbio to collapse sequenced reads into duplex consensus sequences (DCSs) representing the double-stranded original DNA molecules, so both tumor and normal BAM files contain DCSs instead of raw reads. Then, we ran BamSurgeon, VarBen and SafeMut to spike-in mutations into the normal BAM files which have approximately the same depth as the tumor BAM files, generating simulated tumor-like BAM files. Next, we compared the DCS VAF distribution of the real tumor BAM files with respect to the DCS VAF distribution of the simulated tumor-like BAM files.

### Results

We compared the performance of SafeMut with BamSurgeon and VarBen by measuring the similarity between the observed and expected VAF means, VAF variances, and z-score distributions using mean-square errors (MSEs) between the observed and expected means, variances, and distribution quantiles. The observed data were obtained by simulating variants in the normal BAM files and the expected data are directly in the corresponding tumor BAM. Unlike VAF means and variances, z-scores were first sorted, and only then the sorted observed z-scores were compared with the sorted expected z-scores, resulting in the generation z-score Q-Q plots. SafeMut overall performs the best (Figs. 2 and 3). BamSurgeon was affected by a runtime error when simulating variants in the cell-free DNA dataset so that the corresponding results are not available.

**Figure 2:**
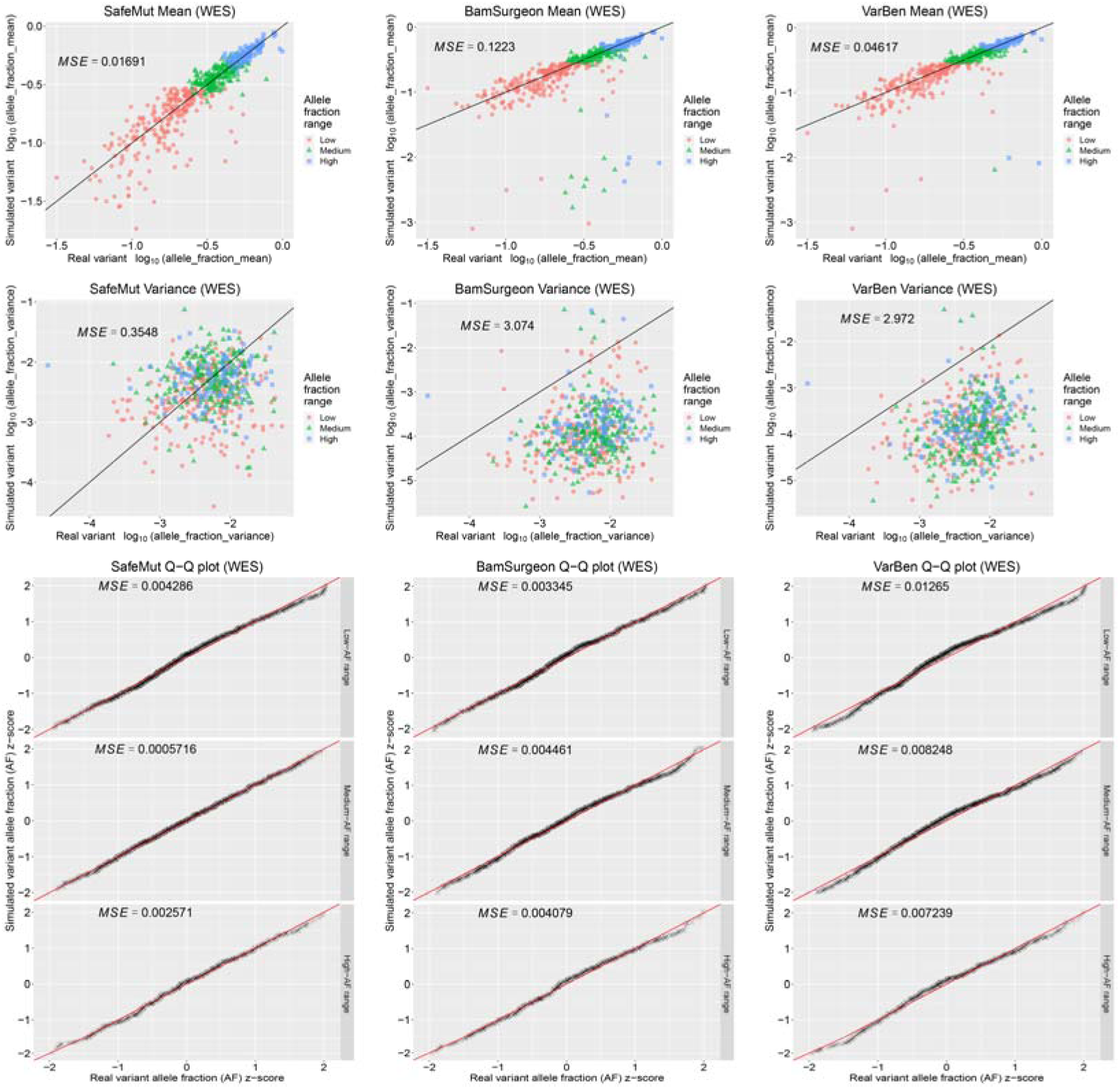
Observed versus expected means (top), variances (mid), and z-scores (bottom) generated by SafeMut (left), BamSurgeon (mid), and VarBen (right) for the HCC1395 whole-exome sequencing (WES) reference standard dataset (accession SRP162370). SafeMut is characterized by the smallest mean-square error (MSE) with respect to the y = x line in all plots except that BamSurgeon is characterized by smaller MSE for the low-AF range z-score Q-Q plot. Low, medium and high AF ranges are delimited by thresholds of 0.3 and 0.5.

**Figure 3:**
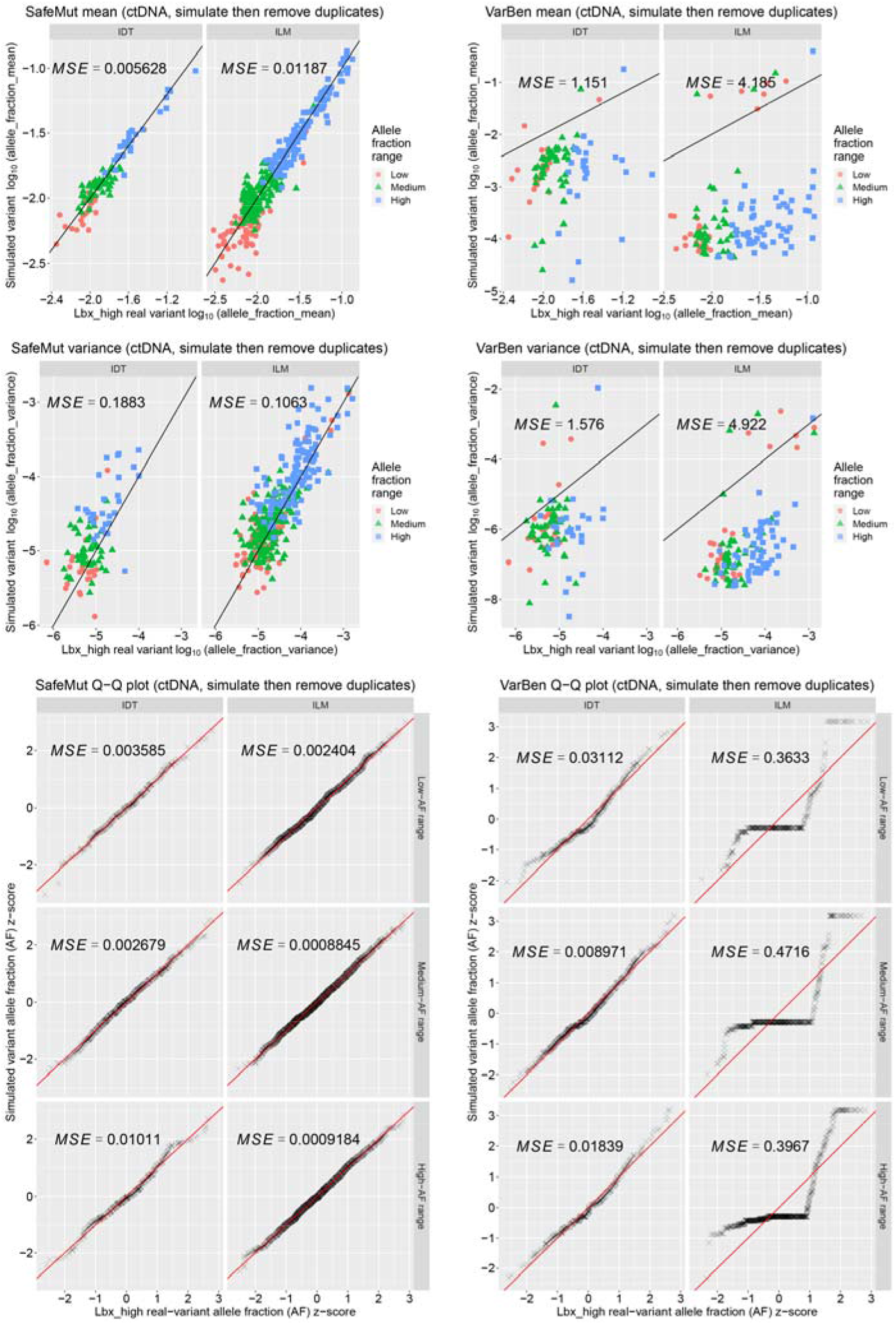
Observed versus expected means (top), variances (mid), and z-scores (bottom) generated by SafeMut (left) and VarBen (right) for the cell-free DNA reference standard dataset (accession SRP296025) evaluated with the simulate-then-consensus strategy. BamSurgeon was affected by runtime error and thus did not generate any result. SafeMut is characterized by the smallest mean-square error (MSE) with respect to the y = x line in all plots. Low, medium and high AF ranges are delimited by thresholds of 0.008 and 0.016.

Overall, SafeMut is characterized by lower MSEs for all combinations of variant simulators (BamSurgeon, VarBen and SafeMut), assay types (whole-exome sequencing (WES), IDT cfDNA assay and Illumina (ILM) cfDNA assay, with both cfDNA assays evaluated with the simulation-then-consensus strategy) and statistical measures (mean, variance, low-AF-variant z-score, medium-AF-variant z-score and high-AF-variant z-score) except that BamSurgeon has lower MSE than SafeMut in the scenario corresponding to the combination of WES and low-AF-variant z-score (Figs. 2 and 3). Moreover, SafeMut is characterized by much smaller MSEs for fitting the linear relationships between observed and expected variances because SafeMut is aware of the AF overdispersion in NGS data. Furthermore, SafeMut strongly outperforms BamSurgeon and VarBen, which are not UMI-aware, on the data generated by UMI-enabled targeted sequencing because SafeMut is UMI-aware (Figs. 2 and 3).

We also evaluated using the cell-free DNA dataset with the consensus-then-simulation strategy (Fig. 4). This strategy enables UMI-unaware simulators to incorporate the effect of the labeling of reads by UMIs in their simulations. However, this strategy is still only a workaround because it discards UMI information such as UMI-family size and UMI-family consensus percent identity. Thus, this strategy disregards UMI-awareness, the advantage that SafeMut has compared with UMI-unaware simulators such as BamSurgeon and VarBen. However, if the consensus-then-simulation strategy is used, then SafeMut still performs the best overall: SafeMut is characterized by the lowest MSEs for all combinations of variant simulators (BamSurgeon, VarBen and SafeMut), assay types (IDT and Illumina cfDNA assays) and statistical measures (mean, variance, low-AF-variant z-score, medium-AF-variant z-score and high-AF-variant z-score) except that VarBen has lower MSEs than SafeMut in the scenarios corresponding to the combinations of both IDT and ILM with mean and that BamSurgeon has lower MSEs than SafeMut in the scenarios corresponding to the combinations of IDT with low-AF-variant z-score and ILM with medium-AF-variant z-score (Fig. 4).

**Figure 4:**
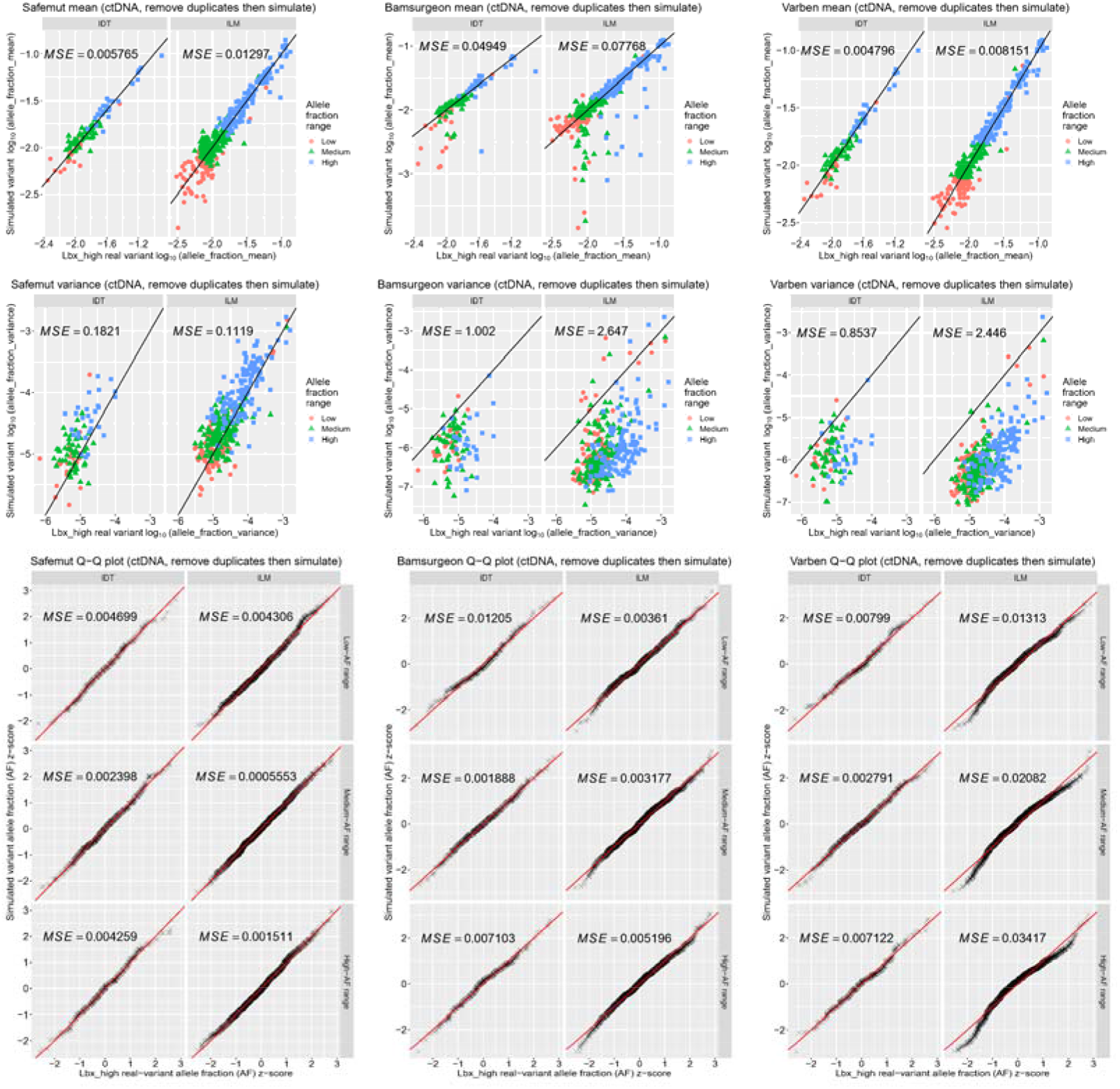
Observed versus expected means (top), variances (mid), and z-scores (bottom) generated by SafeMut (left), BamSurgeon (middle) and VarBen (right) for the cell-free DNA reference standard dataset (accession SRP296025) evaluated with the consensus-then-simulate strategy. Low, medium and high AF ranges are delimited by thresholds of 0.008 and 0.016.

The reference truth AFs, real tumor-data AFs, and simulated tumor-data AFs generated by different bioinformatics tools for each locus are presented at https://doi.org/10.5281/zenodo.7139579 for each sequencing project as detailed results. Importantly, the detailed results show that SNVs and InDels approximately follow the same AF distribution except that simulation by VarBen resulted in over-dispersed InDel AFs but under-dispersed SNV AFs for cfDNA data. Hence, we evaluated SNVs and InDels together instead of evaluating each variant type separately.

The running time (runtime) and random-access memory (RAM) consumed by SafeMut are approximately the same as the respective runtime and RAM consumed by the command “`samtools view -bh`”. And this command is known for its fast runtime and high memory-efficiency in NGS data processing [18]. Therefore, SafeMut is fast and efficient in terms of its usage of computational resources.

## Discussion

The detection of small variants, namely SNVs and InDels, caused by clinically significant mutations are crucial to the diagnosis, prognosis and treatment monitoring of cancer and heritable diseases. Nevertheless, it is labor-intensive and time-consuming to perform wet-lab experiments to obtain the real sequencing data needed for checking the ability of a pipeline to detect such variants. Furthermore, some variants are extremely rare, so a large number of samples have to be collected and sequenced to find the samples characterized by such rare variants, and only then we can check how such variants look like in the sequencing data. Hence, variant simulators become especially important to tackle such problems. Furthermore, the sequencing of cell-free DNA (cfDNA) with molecular barcodes is routinely used in liquid biopsy and minimum residual disease (MRD) detection for cancer patients. As a result, UMI-aware variant simulators are urgently needed, but such simulators have not been developed yet.

We designed and developed SafeMut which simulates small variants (i.e., SNVs and InDels) by editing reads and then *in silico* spike-in the edited reads. SafeMut is applicable to data generated by SafeSeqS and duplex sequencing. Additionally, SafeMut models allele-fraction overdispersion with a specific log-normal distribution. We showed that SafeMut outperforms the state-of-the-art simulators by assessing the similarity between real and simulated variants using two open-access datasets published by the SEQC2 consortium. The variants simulated by SafeMut are extremely similar to real variants in terms of their distributions of allele fractions. Thus, SafeMut is able to provide realistic *in silico* variants which can be used to benchmark the performance for calling these variants. Consequently, SafeMut has numerous applications in bioinformatics. For example, SafeMut can help the user to establish the limit of blank, limit of detection, and limit of quantitation of a rare variant of interest.

SafeMut edits reads in such a way that all variants are in the same haplotype, but sometimes different variants are located in different haplotypes. Hence, the ability to simulate haplotypes may be implemented in a future version of SafeMut. Additionally, SafeMut should be able to work with third-generation sequencing (TGS) data without any modification whatsoever, but further evaluation is needed to assess the performance of SafeMut on TGS.

## Supporting information

Technical details

List of data accessions used in this manuscript.

## Data and code availability

The datasets used to perform the evaluation presented in this work are all accessible to the general public, with accessions listed in (Table S1). The code used to implement SafeMut is publicly available (and free for non-commercial third-party usage) at https://github.com/genetronhealth/safesim with commit 0b4d0f8. The code used to perform the evaluation presented in this work is publicly available at https://github.com/genetronhealth/safesim-eval/ with commit 08c5e8c.

## Acknowledgments

The authors would like to thank Xiaoyue Wang for helping with the coordination between the authors.

## Key Points

- By using UMI-aware generation of variant and by modeling allele-fraction overdispersion with a specific log-normal distribution, we developed SafeMut, an algorithm for spiking-in small variants into existing sequencing data.
- The distribution of the allele fractions (AFs) of the variants that SafeMut spiked-in to the tumor-matched normal samples closely resemble the distribution of the AFs of the real variants directly observed in the corresponding tumor samples.
- Traditionally, sample collections and wet-lab experiments are performed to generate sequencing data of samples characterized by some pre-defined variants. SafeMut alternatively offers a more cost-efficient, less labor-intensive and less time-consuming way to generate such sequencing data by simulating these pre-defined variants *in silico.*

## Reference

1. Huang, W., et al., ART: a next-generation sequencing read simulator. Bioinformatics, 2012. 28(4): p. 593–594.

2. Ewing, A.D., et al., Combining tumor genome simulation with crowdsourcing to benchmark somatic single-nucleotide-variant detection. Nature methods, 2015. 12(7): p. 623–630.

3. Li, Z., et al., VarBen: Generating in silico reference data sets for clinical next-generation sequencing bioinformatics pipeline evaluation. The Journal of Molecular Diagnostics, 2021. 23(3): p. 285–299.

4. Alosaimi, S., et al., A broad survey of DNA sequence data simulation tools. Briefings in functional genomics, 2020. 19(1): p. 49–59.

5. Zhao, M., D. Liu, and H. Qu, Systematic review of next-generation sequencing simulators: computational tools, features and perspectives. Briefings in Functional Genomics, 2016. 16(3): p. 121–128.

6. Cibulskis, K., et al., Sensitive detection of somatic point mutations in impure and heterogeneous cancer samples. Nature biotechnology, 2013. 31(3): p. 213–219.

7. Zack, T.I., et al., Pan-cancer patterns of somatic copy number alteration. Nature genetics, 2013. 45(10): p. 1134–1140.

8. Huang, R.S., et al., Circulating cell-free DNA yield and circulating-tumor DNA quantity from liquid biopsies of 12 139 cancer patients. Clinical Chemistry, 2021. 67(11): p. 1554–1566.

9. Kinde, I., et al., Detection and quantification of rare mutations with massively parallel sequencing. Proceedings of the National Academy of Sciences, 2011. 108(23): p. 9530–9535.

10. Wan, J.C., et al., Liquid biopsies come of age: towards implementation of circulating tumour DNA. Nature Reviews Cancer, 2017. 17(4): p. 223–238.

11. Chin, R.-I., et al., Detection of solid tumor molecular residual disease (MRD) using circulating tumor DNA (ctDNA). Molecular diagnosis & therapy, 2019. 23(3): p. 311–331.

12. Sater, V., et al., UMI-Gen: A UMI-based read simulator for variant calling evaluation in paired-end sequencing NGS libraries. Computational and structural biotechnology journal, 2020. 18: p. 2270–2280.

13. Schmitt, M.W., et al., Detection of ultra-rare mutations by next-generation sequencing. Proceedings of the National Academy of Sciences, 2012. 109(36): p. 14508–14513.

14. Valentine, C.C., et al., Direct quantification of in vivo mutagenesis and carcinogenesis using duplex sequencing. Proceedings of the National Academy of Sciences, 2020. 117(52): p. 33414–33425.

15. Short, N.J., et al., Ultra-accurate Duplex Sequencing for the assessment of pretreatment ABL1 kinase domain mutations in Ph+ ALL. Blood cancer journal, 2020. 10(5): p. 1–9.

16. Ahn, E.H., et al., Detection of ultra-rare mitochondrial mutations in breast stem cells by duplex sequencing. PloS one, 2015. 10(8): p. e0136216.

17. Kim, S., K. Jeong, and V. Bafna, Wessim: a whole-exome sequencing simulator based on in silico exome capture. Bioinformatics, 2013. 29(8): p. 1076–1077.

18. Danecek, P., et al., Twelve years of SAMtools and BCFtools. Gigascience, 2021. 10(2): p. giab008.

19. Zhao, X., et al., Calling small variants using universality with Bayes-factor-adjusted odds ratios. Briefings in Bioinformatics, 2021.

20. Bonfield, J.K., et al., HTSlib: C library for reading/writing high-throughput sequencing data. Gigascience, 2021. 10(2): p. giab007.

21. Jones, L., WG14 N1539 Committee Draft ISO/IEC 9899: 201x. 2010, International Standards Organization.

22. Niu, D., et al., Evaluation of next generation sequencing for detecting HER2 copy number in breast and gastric cancers. Pathology & Oncology Research, 2020. 26(4): p. 2577–2585.

23. Danecek, P., et al., The variant call format and VCFtools. Bioinformatics, 2011. 27(15): p. 2156–2158.

24. Box, G.E., A note on the generation of random normal deviates. Ann. Math. Statist., 1958. 29: p. 610–611.

25. Deveson, I.W., et al., Evaluating the analytical validity of circulating tumor DNA sequencing assays for precision oncology. Nature biotechnology, 2021. 39(9): p. 1115–1128.

26. Fang, L.T., et al., Establishing community reference samples, data and call sets for benchmarking cancer mutation detection using whole-genome sequencing. Nature biotechnology, 2021. 39(9): p. 1151–1160.

27. Li, H., et al., The Sequence Alignment/Map format and SAMtools. Bioinformatics, 2009. 25(16): p. 2078–2079.

28. Li, H., Aligning sequence reads, clone sequences and assembly contigs with BWA-MEM. arXiv preprint arXiv: 1303.3997, 2013.

29. Zerbino, D.R. and E. Birney, Velvet: algorithms for de novo short read assembly using de Bruijn graphs. Genome research, 2008. 18(5): p. 821–829.

30. Slater, G.S.C. and E. Birney, Automated generation of heuristics for biological sequence comparison. BMC bioinformatics, 2005. 6(1): p. 1–11.

31. Jones, W., et al., A verified genomic reference sample for assessing performance of cancer panels detecting small variants of low allele frequency. Genome biology, 2021. 22(1): p. 1–38.

