## Supplementary material for "SafeMut: UMI-aware variant simulator incorporating allele-fraction overdispersion in read editing": Technical details

In the “Algorithm” section of the main text, we used some terms without giving exact definitions, so the definitions are provided here.

The mathematical functions $\mathrm{hash}$ and $\mathrm{hashString}$ are defined as $\_\_ac\_Wang\_hash$ and $\_\_ac\_X31\_hash\_string$ in the khash.h source file from htslib, respectively [1]. The default value for $\mathrm{prngs}$ is $int2randint(13,1)$. The function $int2randint(x, y)$ takes as input an integer $x$ as the PRNG seed, an integer $y$ as the number of iterations, and returns the $y$^th^ random integer generated using this seeded PRNG. In short, the function $int2randint$ returns a PRNG seed from another such seed. The exact implementation of the PRNG used by int2randint is defined by ISO/IEC 9899:201x [2].

1.	Bonfield, J.K., et al., *HTSlib: C library for reading/writing high-throughput sequencing data.* Gigascience, 2021. **10**(2): p. giab007.

2.	Jones, L., *WG14 N1539 Committee Draft ISO/IEC 9899: 201x*. 2010, International Standards Organization.
