## Supplementary material for "SafeMut: UMI-aware variant simulator incorporating allele-fraction overdispersion in read editing": List of data accessions used in this manuscript.

### Supplementary Figures and Tables

| Data | Accessions | |
| --- | --- | --- |
|  | Tumor | Normal |
| WES-data | SRR7890844 | SRR7890878 |
| WES-data | SRR7890876 | SRR7890881 |
| WES-data | SRR7890879 | SRR7890880 |
| WES-data | SRR7890883 | SRR7890966 |
| WES-data | SRR7890918 | SRR7890919 |
| WES-data | SRR7890945 | SRR7890946 |
| ILM-data | SRR13385615 | SRR13385619 |
| ILM-data | SRR13385616 | SRR13385620 |
| ILM-data | SRR13385612 | SRR13385617 |
| ILM-data | SRR13385613 | SRR13385618 |
| ILM-data | SRR13385603 | SRR13385637 |
| ILM-data | SRR13385592 | SRR13385636 |
| ILM-data | SRR13385570 | SRR13385614 |
| ILM-data | SRR13385586 | SRR13385590 |
| ILM-data | SRR13385587 | SRR13385591 |
| ILM-data | SRR13385588 | SRR13385593 |
| ILM-data | SRR13385589 | SRR13385594 |
| ILM-data | SRR13385581 | SRR13385625 |
| IDT-data | SRR13208828 | SRR13208862 |
| IDT-data | SRR13208817 | SRR13208861 |
| IDT-data | SRR13208806 | SRR13208850 |
| IDT-data | SRR13208795 | SRR13208839 |
| IDT-data | SRR13208841 | SRR13208845 |
| IDT-data | SRR13208840 | SRR13208844 |
| IDT-data | SRR13208838 | SRR13208843 |
| IDT-data | SRR13208837 | SRR13208842 |
| IDT-data | SRR13208814 | SRR13208819 |
| IDT-data | SRR13208813 | SRR13208818 |
| IDT-data | SRR13208812 | SRR13208816 |
| IDT-data | SRR13208811 | SRR13208815 |

Table S1. Sequencing data used in our evaluation. WES-data: whole-exome sequencing data. ILM-data: UMI-enabled sequencing data submitted by Illumina. IDT-data: UMI-enabled sequencing data submitted by IDT.
